# Call count surveys as a tool to monitor habitats occupied by Cheer Pheasants (*Catreus wallichii*) in Shimla, Himachal Pradesh

**DOI:** 10.1101/2022.02.17.478413

**Authors:** Samakshi Tiwari, Lakshminarasimha Ranganathan, Sanjeev Kumar, Rajesh Sharma

## Abstract

The vulnerable Cheer Pheasant (*Catreus wallichii*) is found in the western Himalayas. It prefers successional grasslands which are maintained by residents near villages. Therefore, it is prone to anthropogenic pressures which have led to local extinctions and a decline in population. We conducted five call count surveys at dusk and dawn from 15 April – 15 May 2020 in grasslands of Dharbhog panchayat, Shimla, Himachal Pradesh to assess presence of Cheer Pheasants in different habitat types. We used frequency of occurrence (number of encounters / number of visits) to determine presence or absence of Cheer Pheasant in an area of 10.5 km^2^ with ten established call count stations. We encountered Cheer Pheasants at six call count stations and the frequency of occurrence of the species ranged from 0 – 0.60. This survey provides an index of occupancy of the species and can be used to assess the change in habitat occupied by the species with time. The local forest department can also use these results to regulate over grazing and burning of grasslands during the breeding season in areas where we recorded the species.

## Introduction

Cheer Pheasant (*Catreus wallichii*) is a vulnerable galliform found in the western Himalayas ^1^. The sexual dimorphism in the species is not pronounced; with the male being pale grey, barred and having black tail bands, whereas, the female has white streaks on the body, white tail bars and black spots on the breast. The males measure 90–118 cm and female 61–76 cm. Both the sexes have a relatively long tail, crest and red orbital patch. The Cheer Pheasant occurs at an elevation of 1445 – 3050 m from north-eastern Pakistan to central Nepal ^2,3^. It is found in successional habitats which are often maintained by cattle grazing and regular burning of grasslands near villages ^1^. It’s diet predominantly comprises of roots and tubers, and also eats seeds, berries, insects and worms ^1,4^. The species breeds from April to June^3^ during which individuals pair up for the breeding season. They are found in flocks of five or more individuals during the non-breeding season^2^. They dig a shallow nest in grass or at the base of rocks. The female lays clutches of 9-14 eggs ^4^ and incubates for around 26 days ^5^. The parental care is shared by both the sexes.

The species is patchily distributed in the north-western and central Himalaya. Local extinctions from across its distributional range have been reported by various authors ^6–8^. The major threats for the species are grassland burning, habitat degradation, habitat fragmentation and hunting ^9^. Himachal Pradesh is a stronghold of the species distribution. Locally, the species is threatened by habitat fragmentation, nest predation by dogs ^1^, hunting and submergence of key habitats for hydroelectric projects ^10^. The species has been reported to be unable to breed in grasslands prone to burning during the breeding season ^6^.

Other pheasant species endemic to the Himalayas are found in inaccessible high-altitude areas and treacherous terrains. Therefore, they are less threatened by anthropogenic pressures. In contrast, Cheer Pheasants are more prone to such pressures as they are associated with human-use landscapes^11^. Further, the distribution of the species is patchy because of which they are predisposed to local extinction. With the global population of the species ranging between 2,000–2,700 mature individuals ^1^, the species deserves special conservation attention.

Call count surveys are often used to estimate the population of pheasants and to assess their occupancy ^8,11–15^. A study based on visual and call records of the species from all over Himachal Pradesh, estimated that the state holds 1000 pairs of Cheer Pheasant^16^. Another study estimated around to 40 pairs in the Chail wildlife sanctuary ^13^. In 2014, a study used the call count method to report the species from 14 sites across three districts of Himachal Pradesh ^8^. The study also documented local extinction of the species from Jaunaji, Solan district. Conversion of grasslands to agricultural lands and overall reduction in suitable habitat could have resulted in the extinction.

One of the sites assessed during the study (Seri in southern Himachal Pradesh, India) is an important site where the species has been reported historically. A study based on call count survey conducted in March 1988 estimated the population of the species to range from 30 to 35 individuals ^17^. The species was also recorded here in 2014 ^8^ and by different visitors using eBird in 2019, 2020 and 2021^18^. As the species has been recorded here repeatedly, it is likely that it is breeding here. 1 km^2^ area near Seri village was also selected for the reintroduction of the species in 2019^19^.

The remaining wild population of the species is fragmented, declining, and prone to local extinctions. Therefore, it is essential to monitor fragmented small populations of the species with the objective of understanding the factors driving population decline and consequent extinctions at the local level. This can help in developing strategies to reduce anthropogenic pressures and prevent habitat loss. Hence, we conducted surveys to record the presence of Cheer Pheasants in an important distribution area of the species in Himachal Pradesh, India. This should enable subsequent monitoring of local population by the local forest department. This should enable subsequent monitoring of local population by the local forest department.

### Study area

The study was conducted in the grasslands near villages in southern Himachal Pradesh, India (31.04854 - 31.01193°N, 77.27613 - 77.30840°E) covering an area of around 10.5 km^2^. The grasslands are used by residents for fodder collection and cattle grazing. The temperature ranges from −9 °C – 31 °C and the entire study area receives snow in the winter months. The area is surrounded by tributaries of the river Yamuna. The reintroduction of Cheer Pheasants is also ongoing here for which habitat of around 1 km^2^ has been selected for intensive management (20% of the total study area) ^19^.

We laid grids of 500 × 500 m over the study area and carried out a reconnaissance survey for five days. We selected this size as the species has been reported to occupy habitats of similar extent during the breeding season ^6^.

Based on the results of the reconnaissance surveys and local testimony regarding the presence of the species in the grids, we selected ten grids (out of 42) for sampling. Of the ten grids, two grids were within the intensive management area identified as part of the reintroduction project. The terrain at the study site is topographically complex and selection of grids accounted for this by excluding inaccessible areas.

Within each grid we delineated a call count station to record vocal activity of Cheer Pheasant. A call count station consisted of a designated circular area with a 150 m listening radius and stations were positioned at least 500 m to avoid overlap.

## Methods

We mapped the study area, laid grids for sampling using QGIS. We prepared maps for marking the location of calling birds using Google Earth Pro 2018.

Each call count session consisted of 60 minutes of sampling during which vocal activity of Cheer Pheasants was observed. We conducted five call count sessions each at dawn (0645 – 0745 hrs) and dusk (1815 - 1915 hrs) from 15 April - 15 May 2020 (breeding season). We did not conduct the surveys during inclement weather. In total, we invested a total of 47 hours in sampling.

Observers reached the call count stations five minutes prior to each session. No playbacks were used during the sessions. For each call, the observer recorded the number of individuals heard, start time and end time of the call and the number of sighted individuals. We used printed maps of the area to mark locations and then transcribed the same using Google Earth Pro-2018.

We used Microsoft Excel 2007 to calculate the frequency of occurrence at each call count station. For each call count station, we represent the frequency of occurrence (FOO) as the proportion of times at least one calling individual was encountered at a particular station ((number of encounters / number of visits))^20^. We derived duration of calling from by subtracting the start time of call from the end time of each call record.

## Results

We encountered Cheer Pheasants at six call count stations during the five surveys (Table 1). The frequency of occurrence of Cheer Pheasants at four of these call count stations was higher for the evening sessions than the morning sessions.

**Table 1 -.** Frequency of Occurrence ((number of encounters / number of visits)) at call-count stations as observed during the five surveys.

|  | Dawn (0645 – 0745 hrs) |  | Dusk (1815 –1715 hrs) |  | Combined (both the sessions) |  |
| --- | --- | --- | --- | --- | --- | --- |
| Call-count station number<br>(Local name) | Frequency of Occurrence | Sampling time (hours) | Frequency of Occurrence | Sampling time (hours) | Frequency of Occurrence | Sampling time (hours) |
| 1 (Kayad) | 0.20 | 5 | 1 | 5 | 0.60 | 10 |
| 2 (Daraogi) | 0.40 | 5 | 0.60 | 5 | 0.50 | 10 |
| 3 (Kotadu) | 0.20 | 5 | 0.75 | 4 | 0.44 | 9 |
| 4<br>(Seri(Reintroduction Site)) | 0.40 | 5 | 0.40 | 5 | 0.40 | 10 |
| 5 (Shilli Ghati) | 0.20 | 5 | 0.60 | 5 | 0.40 | 10 |
| 6 (Tikri ka Dhaala) | 0.20 | 5 | 0 | 5 | 0.10 | 10 |
| 7 (Undala) | 0 | 5 | 0 | 5 | 0 | 10 |
| 8 (Dhwash) | 0 | 5 | 0 | 5 | 0 | 10 |
| 9 (Bagdra) | 0 | 4 | 0 | 4 | 0 | 8 |
| 10 (Dagan) | 0 | 5 | 0 | 5 | 0 | 10 |

We recorded the highest frequency of occurrence at call count station 1 (Kayad) (1 during the evening session) followed by call count station 2 (Daraogi), 3 (Kotadu), 4 Seri (reintroduction site), 5 (Shilli Ghati) and 6 (Tikri ka Dhaala). No individuals were recorded in four call count stations. We did not record any individuals at call count station 6 (Tikri ka Dhaala) during the evening sessions.

We noted 26 call records in the morning and evening sessions and documented the species visually four times. The duration of calling ranged approximately from 0.5 – 4 minutes (Average −1.33 ± 0.96 minutes).

## Discussion

Based on historical surveys^8,17,18^ and local testimony, we know that a local population of the species has persisted here over the years. Our study period spans over a month and confirms that the species is present in six grids during the breeding season. This can be used as a baseline index to monitor the breeding population of the species over the years.

Increasing anthropogenic pressure has led to local extinctions in some areas ^6–8^. As the population in this area is isolated, it is at a high risk of extinction if not conserved. The species is found in successional habitats which are maintained by cattle grazing and burning of grasses and understory^1^. Therefore, it is essential to maintain such activities while regulating them such that they do not affect the breeding population of the area. Cheer Pheasants at Wacchum in Almora District of Uttar Pradesh are less disturbed during the breeding season as grassland burning and cattle grazing in the area stops by the onset of breeding season ^6^. This allows the shrubs and herbs to grow during the breeding which provides food to the breeding population. Anthropogenic activities in the grasslands we surveyed can be modified to mimic such a timeline.

The Himachal Pradesh Forest Department is already working with residents to monitor reintroduced Cheer Pheasants. With cooperation from the locals, the department has regulated grazing near the soft release pens and has completely stopped burning of grasslands where reintroduced individuals disperse within the intensive management unit. But grasslands which lies outside the intensive management area namely around call count stations 1 (Kayad), 3 (Kotadu), 5 (Shilli Ghati) and 6 (Tikri ka Dhaala) (Figure 1)) were burnt during the breeding season (mainly in June). As these call count stations lie within private grasslands owned by residents of nearby villages, cooperation of the locals is a key in implementing habitat management strategies in these areas.

**Figure 1-.**
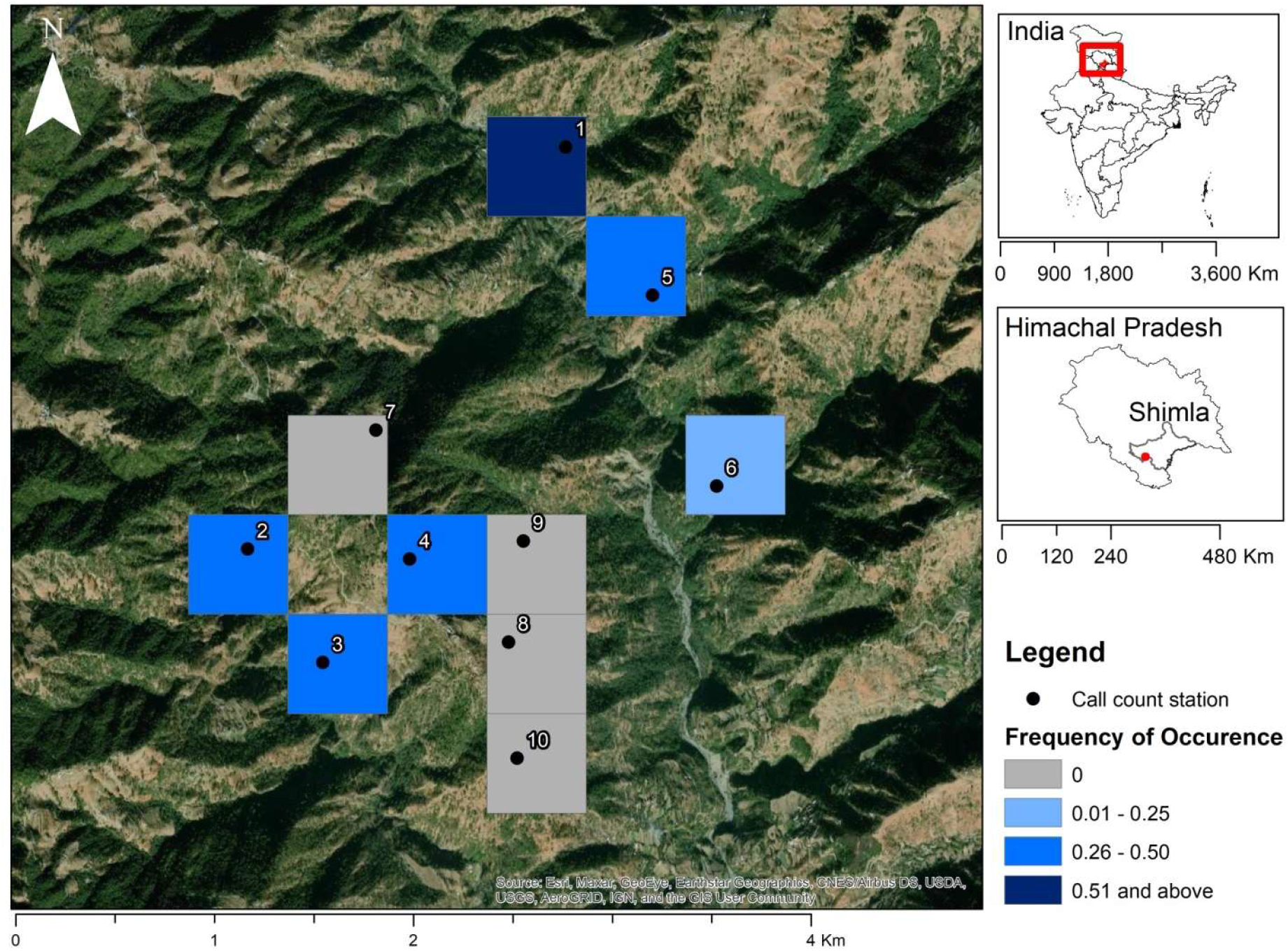
Representation of frequency of occurrence of Cheer Pheasants as recorded in different grids during 10 call count sessions conducted from 15 April - 15 May 2020. Call count stations-1-Kayad, 2-Daraogi, 3-Kotadu, 4-Seri (Reintroduction Site), 5- Shilli Ghati, 6- Tikri ka Dhaala, 7- Undala, 8 - Dhwash, 9- Bagdra and 10- Dagan. All state and boundaries are as per Survey of India.

We did not record the species at call count stations 8 (Dhwash) and 10 (Dagain) (Figure 1). But these lie in continuation with call count station 3 (Kotadu) where Cheer Pheasants were recorded with a frequency of occurrence of 0.44. This finding is supported by previous observations ^4,21^. Other studies too found that the Cheer Pheasant is very patchily distributed and it may be absent from areas which apparently look like other habitats where it has been recorded. Based on this survey, information on potential areas for reintroductions in future may be identified. Further studies of the vegetation structure, predation pressure and anthropogenic disturbances of habitats occupied by the species can be carried out to understand the Cheer Pheasant’s habitat preferences.

This is the first report of frequency of occurrence of the species from different regions of Dharbhog panchayat. By following the changes in frequency of occurrence over the years, an index of trend in change in extent of habitat inhabited by the species can be estimated in the future. This report is based only on the surveys conducted during the breeding season. As Cheer Pheasants are vocally active throughout the year (except during the immediate post-breeding season) ^12^, similar surveys can be conducted in different seasons to understand how the extent of habitats occupied changes throughout the year. The effort can also be extended to a larger study area and used to estimate the population of the species in larger panchayats and districts.

## Acknowledgements

We are grateful to all members of the Cheer Pheasant reintroduction team, Himachal Pradesh Forest Department, specifically the Wildlife Wing and residents (owners) for allowing us to execute the study. We also thank Pr. CCF Wildlife Shimla and CEO HPZCBS for facilitating the study. We are grateful to residents Vikas Thakur and Sunil Sharma for helping in data collection. We thank Puja Sharma, Tim Inskipp, Rajah Jayapal and staff of SACON and WII library for providing reference materials and Shailja Mamgain for helping in representing the study area.

## Notes

### Competing Interest Statement

The authors have declared no competing interest.

